# *Dcaf17* knockdown integrates mitochondrial-lysosomal dysfunction with ferroptosis, NLRP3 inflammasome signaling and necroptosis

**DOI:** 10.1101/2025.06.04.657829

**Authors:** Nadya Alyacoub, Falah Almohanna, Alanoud Alqassem, Salma Awad Mahmoud, Amer Almzroua, Abdullah mohamed Assiri

## Abstract

DCAF17 (DDB1-and CUL4-associated factor 17) plays a vital role in spermatogenesis, influencing post-meiotic sperm development. Its dysfunction leads to male infertility, as demonstrated in our previous study on *Dcaf17* knockout mice, which revealed impaired sperm morphogenesis, increased numbers of dysmorphic and immotile sperm, and complete male infertility. Mutations in the human *Dcaf17* gene have been implicated as causative factors for Woodhouse-Sakati Syndrome (WSS), a disorder characterized by hypogonadism and infertility, highlighting the gene’s essential role in male reproductive function. Despite these observations, the molecular mechanisms underlying DCAF17 function remain poorly defined. Here, we employed the GC1 spermatogonia cell line to investigate the role of DCAF17 in germ cell viability and homeostasis. Lentiviral shRNA-mediated knockdown of *Dcaf17* significantly decreased cell viability and increased cell death. Functional studies demonstrated a notable decline in mitochondrial membrane potential and mass, coinciding with increased oxidative phosphorylation (OXPHOS) protein expression. Unexpectedly, both mitochondrial and cytosolic reactive oxygen species (ROS) levels were reduced following *Dcaf17* silencing. Further investigation uncovered profound lysosomal damage, evidenced by accumulation of SQSTM1/p62 and loss of acidity. Transcriptional and translational profiling indicated downregulation of key iron transport and antioxidant genes (xCT, transferrin), while levels of transferrin receptor were elevated. Silencing of *Dcaf17* triggered ferroptosis, as evidenced by elevated Fe²⁺, reduced glutathione (GSH) and increased lipid peroxidation (MDA). In parallel, inflammatory and necroptotic pathways were activated, as indicated by upregulation of MLKL, NLRP3, cleaved IL-1β, and phosphorylated JNK1/2, along with increased LDH release and reduced Bcl-xL and Bcl-2. These molecular signatures were recapitulated in lymphoblastoid cell lines from WSS patients with a pathogenic *Dcaf17* mutation, which exhibited mitochondrial and lysosomal dysfunction, disrupted iron homeostasis, and oxidative stress. Similar alterations were observed in testis tissue and sperm from *Dcaf17* knockout mice, including increased Fe²⁺ and MDA levels, reduced GSH, and impaired mitochondrial function in sperm. Collectively, our findings establish DCAF17 as a key regulator of mitochondrial integrity, lysosomal function, iron metabolism, and regulated cell death pathways in spermatogonia. This study provides the first mechanistic insight into how *Dcaf17* deficiency contributes to male infertility.

## Introduction

Cellular homeostasis is governed by the coordinated functions of mitochondria and lysosomes, two critical organelles involved in energy production, metabolic regulation, and adaptation to stress [1]. Beyond their canonical roles, both organelles participate in signal transduction, cell fate determination, and quality control mechanisms critical for tissue-specific functions. Lysosomes through vesicular trafficking and degradation via hubs containing acid hydrolases, recycle macromolecules and modulate cellular signaling, autophagy, nutrient sensing, and apoptosis [2, 3]. Lysosomal dysfunction impairs autophagy, leading to proteotoxic stress and organelle accumulation, which contribute to neurodegeneration, metabolic disorders, and immune dysregulation [4]. The male reproductive system depends on effective lysosomal activities to safeguard normal spermatogenesis, sperm development, and fertilization capability. Sertoli cell lysosomes clear apoptotic germ cells and maintain the seminiferous epithelium, while contributing to germ cell support via the secretion of regulatory factors [5–9]. In the epididymis, lysosomes modulate sperm surface remodeling, and during fertilization, lysosomal enzymes enable the acrosome reaction necessary for oocyte penetration [10, 11]. Mitochondria are equally integral to testicular function. Their function is for ATP synthesis, cellular energy balance, regulating calcium flux, redox homeostasis and apoptosis signaling. Mitochondrial dynamics, including fusion and fission are particularly critical during spermatogenesis, influencing stem cell fate and differentiation [12–15]. Mitochondrial remodeling during spermiogenesis ensures proper sperm structure and function, and their localization in the sperm midpiece provides localized energy for motility. [14, 16, 17]. Impaired mitochondria function often manifested as impaired ATP production, elevated reactive oxygen species (ROS) levels, and disrupted calcium homeostasis, can lead to the activate of various forms of programmed cell death (PCD), such as apoptosis, necroptosis, and ferroptosis [18]. These death pathways are increasingly recognized as modulators of spermatogenic integrity and male fertility. Ferroptosis a lipid peroxidation–driven, iron-dependent form of PCD, is active in spermatogenic cells, where it impairs meiosis and spermatogenesis [19]. Environmental and chemical toxins (e.g., cadmium, PM2.5), aging, and genetic predispositions aggravate ferroptosis by promoting iron overload and oxidative stress [20–26]. Similarly, necroptosis, mediated by RIPK1, RIPK3, and MLKL, has been implicated in age-related infertility [27]. Moreover, functional crosstalk between lysosomes and mitochondria is increasingly recognized as critical for maintaining organelle quality control, metabolic coordination, and stress response [28]. Disruption of this bidirectional communication can amplify stress signals and to disease pathogenesis, including cancer, cardiovascular disease, and neurodegeneration [29–31]. Notably, disruptions in mitochondrial–lysosomal crosstalk have been recently associated with decreased sperm motility and membrane integrity in conditions such as bilateral varicocele [32]. Nevertheless, the upstream molecular regulators orchestrating these organelle dysfunctions in the testis remain poorly understood.

The testis-enriched E3 ligase substrate receptor DCAF17, a component of the CUL4– DDB1 complex, is crucial for male germ cell maturation. Mutations in *Dcaf17* cause Woodhouse-Sakati Syndrome, a recessive disorder in which male infertility and hypogonadism are prominent features [33], DCAF17 is highly expressed in the testis, with relatively low expression observed in other tissues; suggesting a specialized role in male reproductive biology [34]. Knockout mouse models have demonstrated that *Dcaf17* is essential for normal spermatogenesis [34, 35]. Despite its established importance, the molecular mechanisms by which DCAF17 regulates spermatogenesis remain poorly understood.

In this study, we investigated the role of DCAF17, using RNAi-mediated knockdown in GC1 spg spermatogonial cells, complemented by validation in WSS patient-derived lymphoblastoid cells (LCLs) harboring a homozygous *Dcaf17* mutation. We demonstrate that *Dcaf17* deficiency disrupts mitochondrial function, impairs lysosomal integrity, and alters iron metabolism, culminating in increased susceptibility to cell death. Our findings characterize DCAF17 as a novel upstream effector connecting organelle disruption with regulated necrosis in the testis. Our findings offer, to the best of our knowledge, for the first-time insights into the molecular mechanisms that underpin not only the spermatogenic defects and male infertility associated with *Dcaf17* knockout, but also pave the way for a preliminary comprehension of the pathogenesis WSS.

## Methods

### Cell culture

Patient-derived lymphoblastoid cell lines (LCLs) carrying a pathogenic *Dcaf17* variant— three independent patient isolates—and one control line were kindly provided by Dr Fowzan Alkuraya (King Faisal Specialist Hospital & Research Centre, Riyadh, Saudi Arabia). All LCL-based experiments were performed with this fixed panel of four lines (n = 1 control, n = 3 mutant). LCLs were expanded in RPMI-1640 medium supplemented with 15 % (v/v) fetal bovine serum, 2 mM L-glutamine, and 1 % (v/v) penicillin-streptomycin, and propagated in six-well plates and subcultured at a ratio of 1:3 every other day. HEK 293T/17 cells (CRL-11268, ATCC), utilized for lentivirus production, and murine testicular germ cell line GC-1 spg (CRL-2053, ATCC) were maintained in high-glucose Dulbecco’s modified Eagle medium containing 10 % (v/v) fetal bovine serum and 1 % (v/v) penicillin-streptomycin. All cell cultures were maintained at a 37 °C incubator with a 95% humidified air atmosphere and 5 % CO₂ atmosphere. All functional assays utilizing GC-1 spg cells were carried out with three GC-1 spg–derived cell populations: (i) GC-1 spg cells stably expressing a non-targeting scramble shRNA (control), (ii) GC-1 spg cells expressing *Dcaf17*-targeting shRNA sequence 1, and (iii) GC-1 spg cells expressing *Dcaf17*-targeting shRNA sequence. All cultures were kept at 37 °C in a humidified atmosphere of 95 % air and 5 % CO₂ throughout the study.

### Generation of shRNA lentivirus and transduction of GC-1 spg cells

Small interfering RNA (siRNA) sequences targeting the mouse *Dcaf17* gene (RefSeq: NM\_001165980.1) were designed using the siRNA design tool provided by Dharmacon. The top two siRNA candidates were synthesized, annealed, and cloned downstream of the U6 promoter into the third-generation lentiviral vector pLL3.7 (Addgene) using HpaI and XhoI restriction sites (New England Biolabs). A non-targeting scrambled shRNA construct was generated using the same cloning strategy to serve as a negative control. The resulting constructs were used to produce VSV-G pseudotyped, replication-deficient lentiviral particles following the protocol described by AlYacoub et al. [36]. For viral transduction, GC-1 spg cells were resuspended in 2 mL of virus-containing supernatant (SN) at a density of 8 × 10⁵ cells per 2ml SN. Polybrene was added to a final concentration of 8 µg/mL and the virus-cell suspension was centrifuged at 2,000 rpm for 1 hour at room temperature. Following centrifugation, cells were plated directly in the viral supernatant, which was replaced with fresh complete medium after 12 hours. Transduction efficiency was evaluated by measuring GFP fluorescence-encoded by the pLL3.7 vector-using flow cytometry.

### Determination of Cell Viability cell death and LDH release

Cell viability was quantified with the colorimetric WST-1 assay (Cell Proliferation Reagent WST-1, Roche), which is based on the reduction of the tetrazolium salt WST-1 to soluble formazan by mitochondrial dehydrogenases in metabolically active cells. Cells were seeded at 6 × 10^4 cells per well in 96-well plates containing complete growth medium. Following a 24-hour incubation period, WST-1 reagent was added directly to the cultures and subsequently incubated for 30 minutes at 37 °C for 30 minutes. The absorbance of the formazan product was measured at 450 nm using a microplate reader. Cell viability was expressed as the percentage of viable cells relative to that observed in scramble shRNA-expressing cells.

To quantify cell death induction, GC-1 spg cells were seeded in a 6-well plate at 8× 104 cells/well in 2ml of complete culture medium and subsequently incubated for approximately 16 hours. Subsequently, the cells were collected and apoptosis was quantified using the Cell Death Detection ELISA kit (Cell Death Detection ELISA, Roche) following the manufacturer’s instructions.

Necroptotic activity was assessed with the LDH (lactate dehydrogenase) release assay, which measures cell membrane integrity and cytotoxicity. Culture supernatants (20 µL) were harvested from the same experimental wells used in the preceding cell death assay and LDH was quantified using the LDH Detection Kit (Cytotoxicity Detection Kit, Roche) according to the manufacturer’s instructions. Both LDH release and cell-death values were normalized to scramble-shRNA cells and expressed as fold change.

### Cell cycle analysis

Cell cycle distribution was assessed using propidium iodide (PI) staining, following the protocol established by Nicoletti et. al. [37] Cells were seeded in 6-well plates at 8 × 10⁴ cells per well in 2 mL of complete medium and incubated for 18 hours. After incubation, cells were harvested, washed with cold phosphate-buffered saline (PBS), and incubated for 1 hour at 4 °C in the dark in a hypotonic staining solution containing 40 µg/mL PI, 0.1% sodium citrate, and 0.1% Triton X-100. DNA content was then analyzed using a BD Accuri-C6 flow cytometer (BD Biosciences), with PI fluorescence detected in the FL2-A channel to determine cell cycle distribution, including the G1 population. As this PI-based protocol does not preserve GFP fluorescence, the standard gating strategy used for isolating GFP-positive cells could not be applied. To ensure that observed differences in cell cycle profiles between scramble and *Dcaf17*-shRNA–expressing cells were attributable to *Dcaf17* knockdown, a small aliquot of each sample was taken prior to PI staining to assess the percentage of GFP-positive cells by flow cytometry. Only samples in which > 50% of the population was GFP-positive were included in the final cell cycle analysis.

### Assessment of mitochondrial membrane potential and mitochondrial mass

To assess mitochondrial membrane potential (ΔΨm), cells were collected washed with PBS and then stained with tetramethylrhodamine methyl ester (TMRM+, Sigma-Aldrich). Following trypsinization, cells were incubated with 5 µM TMRM+ for 20 minutes at 37 °C in the dark. Following two washes with PBS supplemented with 1% BSA, cells were resuspended in staining buffer and analyzed using the Accuri-C6 flow cytometer. For this and all subsequent flow cytometry-based assays, a gating strategy was applied that relied on a non-transfected, unstained double-negative control. Initial gating was performed to exclude debris and doublets by selecting single cells based on FSC-H vs. FSC-A parameters. GFP-positive cells were then gated to exclude untransfected populations, ensuring that only transfected cells were analyzed. Within the GFP-positive gate, TMRM fluorescence was assessed, with unstained GFP-positive controls used to establish baseline fluorescence and apply spectral compensation where needed to correct for overlap between GFP and TMRM signals. Only GFP-positive cells were included in the analysis, as these represent the population in which *Dcaf17* was knocked down, and thus reflect the specific biological effects under investigation.

Mitochondrial mass was assessed by staining cells with 1 µM MitoTracker Red CMXRos (#M46752; Thermofisher scientific). GC-1 spg cells were incubated with the probe for 30 minutes, and LCLs for 1 hour, at 37 °C in PBS with 1% BSA. Following two washes with cold PBS, cells were analyzed by flow cytometry using the same gating strategy employed for TMRM staining.

### Assessment of ROS generation

The intracellular total ROS levels were assessed using the CellROX Deep Red probe (#C10422; Thermofisher scientific), while mitochondrial superoxide production was measured using the MitoSOX Red probe (#M36008; Thermofisher scientific). For both assays, cells were harvested, washed with PBS, and incubated with either 5 µM CellROX Deep Red or 5 µM MitoSOX Red for 20 minutes at 37 °C in the dark. Following two washes with PBS, stained cells were resuspended in buffer and analyzed by flow cytometry. The gating and spectral compensation strategy used was identical to that applied for TMRM analysis to focus the analysis on the GFP-positive population.

### Quantification of mtDNA levels by qPCR

Total genomic DNA was extracted using the Puregene Core A Kit (QIAGEN), following the manufacturer’s protocol, and quantified using a NanoDrop 2000 spectrophotometer (Thermofisher scientific). Mitochondrial DNA (mtDNA) level was determined by qPCR using the 7500 Fast Real-Time PCR System (Applied Biosystems) employing the Hk2 primers for nuclear DNA and ND1 and ND5 for mtDNA. Each qPCR reaction was performed using 30 ng of template DNA and the PowerUp™ SYBR™ Green Master Mix (#A25743; Thermofisher scientific), with the thermal cycling conditions set according to the manufacturer’s recommendations. The mtDNA content was calculated and expressed as mitochondrial genome copies per nuclear genome using the formula 2^ΔCt. Only samples in which > 50% of the population was GFP-positive were included in the final cell cycle analysis.

### Detection of lysosomal membrane permeabilization

To evaluate the loss of lysosomal acidity as an indicator of lysosomal membrane integrity, cells were stained with either LysoTracker-Deep Red (#L12492, Thermofisher scientific) or Acridine orange (AO, #A1301; Thermofisher scientific) and analyzed by flow cytometry. LysoTracker is a cell-permeable, protonated fluorescent dye that preferentially accumulates in acidic lysosomes, while AO is a weak base that diffuses into cells and accumulates in acidic compartments, emitting red fluorescence in lysosomes and green fluorescence in the cytosol. Following PBS washes, cells were incubated with 100 nM LysoTracker for 1 hour or 200 nM AO for 20 minutes at 37 °C in the dark. After staining, cells were washed twice with cold PBS containing 1% BSA and resuspended in staining buffer for flow cytometric analysis employing the previously described gating strategy. Given the spectral overlap with GFP, only the red fluorescence signal from AO (AO-Red) was used for analysis, excluding the green channel.

### Assessment of oxidative stress and free iron content

Oxidative stress and antioxidant capacity were evaluated by quantifying malondialdehyde (MDA), an indicator of lipid peroxidation, and total glutathione (GSH) levels in both cultured cells and testicular tissue. MDA levels were measured using a colorimetric thiobarbituric acid reactive substances (TBARS, Cayman) assay, in which MDA reacts with thiobarbituric acid (TBA) to form a chromogenic adduct detectable by spectrophotometry. Total GSH content was determined using a colorimetric assay kit (Glutathione Assay Kit, Cayman) based on an enzymatic recycling method involving glutathione reductase, which enables accurate quantification of reduced glutathione. In addition, intracellular iron levels—including total iron, ferrous (Fe²⁺), and ferric (Fe³⁺) forms—were quantified using a colorimetric iron assay kit (Abcam). All assays were performed strictly in accordance with the protocols outlined by the manufacturer. Final concentrations of MDA, GSH, and iron were calculated according to each kit’s standards and normalized to the total protein content of each sample (expressed per mg protein). Only samples in which >50% of the population was GFP-positive were included in the final cell cycle analysis.

### RNA isolation, cDNA synthesis and Real-Time PCR analysis

Total RNA was isolated from cultured cells using TRIzol® Reagent (Thermofisher scientific), following the manufacturer’s protocol. RNA concentration and purity were assessed using a NanoDrop 2000 spectrophotometer (Thermofisher scientific). For each sample, 2.5 µg of total RNA was used for cDNA synthesis. Reverse transcription was performed using the SuperScript™ IV VILO™ Master Mix (Thermofisher scientific), which includes ezDNase for removal of contaminating genomic DNA, following the supplier’s instructions. Quantitative real-time PCR (qRT-PCR) was carried out on a 7500 Fast Real-Time PCR System (Applied Biosystems, CA, USA) using PowerUp™ SYBR® Green Master Mix and gene-specific primers. Each reaction contained 3 µL of 1:8 diluted cDNA, primers at a final concentration of 5 µM, and a total reaction volume of 20 µL. PCR cycling conditions followed the manufacturer’s recommendations for the SYBR Green mix, with primer-specific optimization of the annealing temperature. Primer specificity was confirmed by melting curve analysis. All qRT-PCR reactions were performed in technical duplicates for each of the biological replicates. Relative gene expression levels were calculated using the 2^−ΔΔCt method, with B2M or 18sRNA serving as internal reference genes. Primers were designed using the online qPCR Primer Design Tool provided by Integrated DNA Technologies (https://www.idtdna.com).

### Protein extraction and western blot analysis

GC-1 spg cells were lysed by sonication in a high-salt lysis buffer composed of 20 mM HEPES (pH 7.5), 0.65 M NaCl, 1 mM EDTA, and 0.34 M sucrose, supplemented with protease & phosphatase inhibitor cocktail (Thermo Fisher Scientific). Cell lysates were clarified by centrifugation at 12,000 rpm for 15 minutes at 4 °C, and the resulting supernatants were collected. Protein concentrations were determined using the BCA Protein Assay Kit (Thermofisher scientific) following the manufacturer’s protocol. For immunoblotting, equal amounts of total protein (20–30 µg) were resolved by SDS-PAGE and transferred onto 0.45 µm nitrocellulose membranes (GE Healthcare, Amersham). Membranes were blocked for 1 hour at room temperature with 5% non-fat milk in TBST, followed by overnight incubation at 4 °C with the appropriate primary antibodies at the indicated dilutions: ERK1/2 (Cell Signaling Technology [CST], 1:1000), phospho-ERK1/2 (CST, 1:1000), Bcl-xL (CST, 1:300), Bcl-2 (Novus, 1:200), β-Actin (CST, 1:1000), MLKL (Proteintech, 1:600), JNK1/2 (Santa Cruz Biotechnology, 1:800), phospho-JNK1/2 (Santa Cruz, 1:200), OxPhos Rodent WB Antibody Cocktail (Thermofisher scientific, 1:1000), Transferrin (TF; Thermofisher scientific, 1:300), Transferrin Receptor (TFRC; Thermofisher scientific, 1:700), Ferritin Heavy Chain (FTH1) and Light Chain (FTL1) (Thermofisher scientific, 1:200), and xCT (GeneTex, 1:300). After three washes with TBST membranes were incubated for 90 minutes at room temperature with respective horseradish peroxidase-conjugated (HRP)-conjugated secondary antibodies (CST, 1:3000). Blots were washed three additional times and developed using the Pierce ECL Western Blotting Substrate (#34577; Thermofisher scientific). Chemiluminescence signals were visualized and documented using the Amersham 680 Imaging System (GE Healthcare). For Western blot analysis, GFP-positive cells were further enriched by fluorescence-based cell sorting to increase sample purity prior to protein extraction. Due to technical limitations, the typical post-sorting purity achieved was approximately 70%.

### Animal Handling, Tissue Collection, and Sperm Isolation

Eight-week-old male mice carrying a targeted deletion of the *Dcaf17 gene* (knockout; KO) and age-matched wild-type (WT) C57BL/6J controls were obtained as previously described [34]. All animals were obtained from the Laboratory Animal Services Unit at King Faisal Specialist Hospital and Research Centre (KFSHRC), Riyadh, Saudi Arabia.

All procedures involving animals were conducted in compliance with ethical regulations and approved by the Animal Care and Use Committee at KFSHRC (Project RAC#2160019). Mice were housed under standard conditions with a 12-hour light/dark cycle and had free access to rodent chow and water throughout the study. Euthanasia and tissue collection were performed in accordance with the Institutional Animal Care and Use Committee (IACUC) guidelines and the National Research Council’s Guide for the Care and Use of Laboratory Animals. For biochemical assays involving the testis, samples were rapidly snap-frozen in liquid nitrogen immediately after dissection and stored at −80 °C until further processing.

For sperm staining assays, both testis and epididymis were collected. Sperm from WT and *Dcaf17*-KO mice were isolated from cauda epididymides as previously described [38]. Briefly, the cauda epididymis was immediately placed in pre-warmed M2 medium (Sigma-Aldrich) in a 60 mm petri dish, pierced, and incubated at 37 °C for approximately 30 minutes to allow motile sperm to swim out. The sperm-containing medium was then collected and centrifuged at 500 × g for 10 minutes at room temperature. The resulting sperm pellet was resuspended in warm PBS containing a protease inhibitor cocktail (Thermofisher scientific). After counting, equal numbers of sperm were aliquoted into separate tubes for staining with either MitoSOX Red or TMRM probes, following the staining protocols described above.

## Statistical Analysis

Unless otherwise specified in the Figure legends, all data presented are representative of at least three independent experiments. Results are expressed as the mean ± standard error of the mean (SEM). Statistical significance was calculated using a two-tailed Student’s t-test. Significance levels are denoted in the Figures as follows: p ≤ 0.05 (*), p ≤ 0.01 (**), p ≤ 0.001 (***), and p ≤ 0.0001 (****).

## Results

### *Dcaf17* knock-down compromises viability and induces cell death in GC-1 spg spermatogonial cells

Previously, we have reported that systemic loss of *Dcaf17* has been indicated with increased germ cell apoptosis with varying degrees depending on the specific stages of germ cell development [34]. However, the mechanisms by which DCAF17 regulates spermatogenesis remain unclear. To investigate this, we used the GC-1 spg mouse spermatogonial cell line as an in vitro model to examine DCAF17 function in early spermatogenesis. Stable knockdown of *Dcaf17* was achieved using lentiviral-mediated delivery of two shRNA sequences targeting *Dcaf17*. Real-time PCR analysis confirmed that both antisense oligonucleotides efficiently silenced *Dcaf17* expression (Fig. 1a). Thus, lentivirus particles carrying these two antisense sequences were in all subsequent experiments.

**Figure 1.**
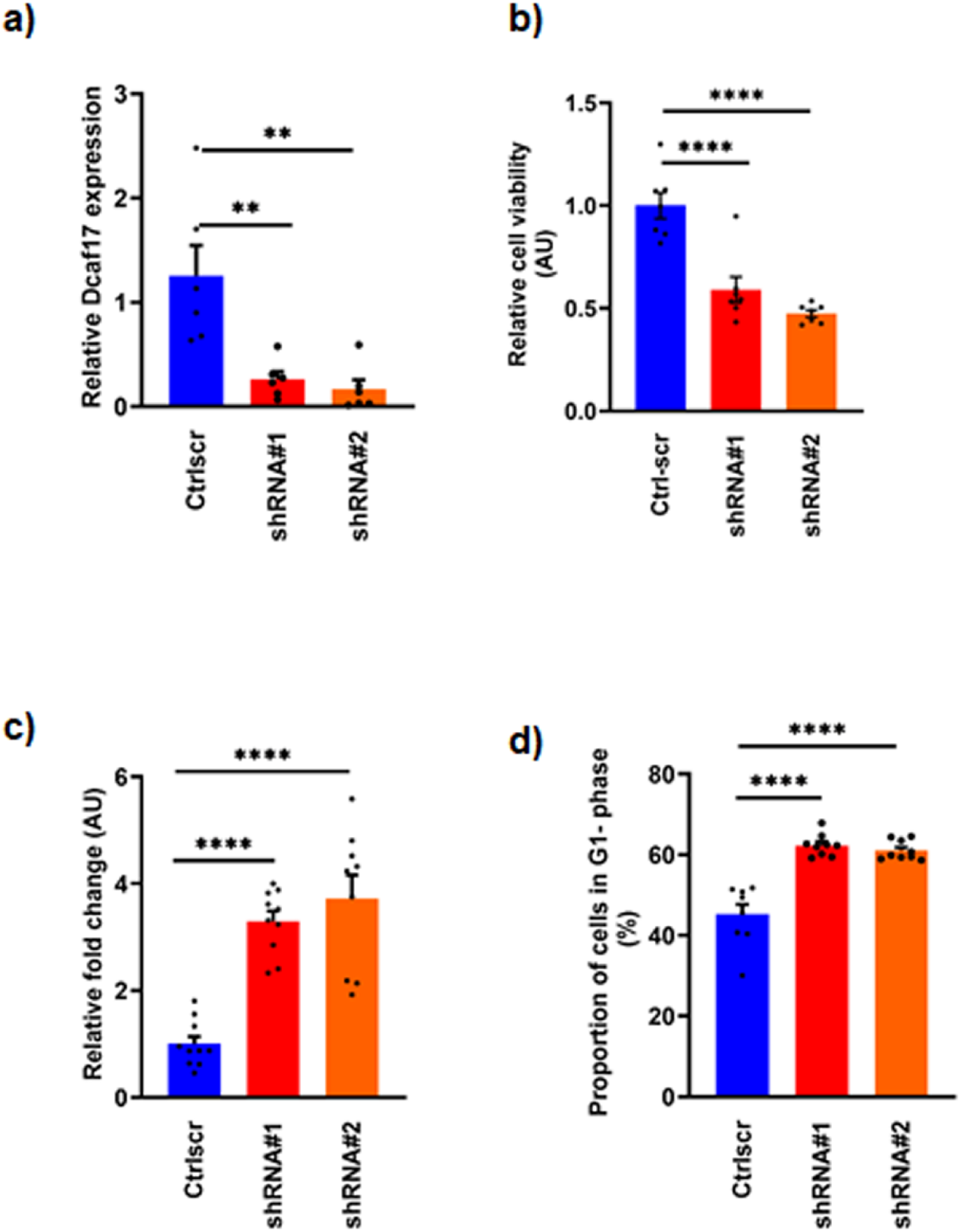
***Dcaf17* knockdown reduces viability, promotes apoptosis, and induces G1 arrest in GC-1 spg spermatogonia.** (a) GC-1 spg cells stably expressing scrambled shRNA (scr) or two independent *Dcaf17*-targeting shRNAs (shRNA#1 and shRNA#2) were used for all analyses. (a) Steady-state *Dcaf17* mRNA levels were quantified by RT– qPCR and normalized to 18S rRNA. Values are presented relative to scr. (b) Cell viability was assessed using the WST-1 assay and expressed as fold change relative to scr (set to 1). (c) Apoptosis was quantified using a Cell Death Detection ELISA. (d) Cell cycle distribution was assessed by flow cytometry following propidium iodide staining; G1 phase percentage is shown. Data represent mean ± s.e.m. from at least three biologically independent experiments. Statistical significance was calculated using two-tailed unpaired Student’s t-test. p ≤ 0.05 (*), p ≤ 0.01 (**), p ≤ 0.001 (***), and p ≤ 0.0001 (****). Data are shown as arbitrary units (AU).

Next we examined the impact of *Dcaf17* knockdown on cell survival, using WST assay, and Cell death assay to analyze cell proliferation, and apoptosis, respectively. *Dcaf17* knockdown led to a significant reduction in cell proliferation and a corresponding increase in apoptosis compared to cells expressing scrambled shRNA (Fig. 1b-c). Flow cytometry analysis of DNA content revealed an increase in the proportion of cells in the G1 phase, indicating G1-phase cell cycle arrest following *Dcaf17* knockdown (Fig.1d). Collectively, these results indicate that DCAF17 is essential for cell survival and progression through the cell cycle in GC-1 spg cells.

### *Dcaf17* silencing impairs mitochondrial function and reduces mitochondrial mass in GC-1 spg cells

Mitochondria play a crucial role in numerous cellular processes, including energy production and cell death, and are essential for the development and functionality of germ cells in testicular tissue. Alterations in mitochondrial physiology have detrimental effects on sperm quality and have been linked to male infertility. Furthermore, we have previously reported that *Dcaf17*^−/−^mutant sperm display morphological abnormalities [34]. To evaluate mitochondrial function in *Dcaf17*-silenced GC-1 spg cells, we assessed mitochondrial membrane potential (ΔΨm) using TMRE (ethyl ester, perchlorate) staining and flow cytometry. A marked reduction in TMRE fluorescence was observed in *Dcaf17* knockdown cells compared to controls, indicating mitochondrial depolarization (Fig. 2a). Having established that the *Dcaf17* knockdown negatively affects mitochondrial membrane potential, we next asked whether basal levels of oxidative phosphorylation (OXPHOS) were affected. Western blot assessment of subunits of the OXPHOS complexes showed a strong increase in the expression of all mitochondrial ETC complexes except for complex I in *Dcaf17*-knockdown compared to scramble cells (Fig. 2b). Mitochondria are considered the major cellular compartment contributing to the production of reactive oxygen species (ROS) in mammalian cells. The imbalance between mtROS production and removal affects negatively several cellular components such as proteins, lipids, and DNA [39]. Therefore, we investigated whether *Dcaf17* knockdown correlates with altered ROS production. To this end we employed the two ROS probes, the MitoSoxRed and the CellRox Deep Red to determine mitochondrial ROS content and the overall ROS content respectively. Flow cytometry revealed a significant decrease in both mitochondrial and overall ROS levels in *Dcaf17*-silenced cells (Fig. 2c&d). A decrease in mitochondrial ROS could reflect a decrease of mitochondria mass in relation to the cell volume. Therefore, we examined whether these observed altered mitochondrial function are accompanied by changes in mitochondrial mass. demonstrated a significant decrease in fluorescence intensity in *Dcaf17*-deficient cells, indicating reduced mitochondrial mass (Fig. 2e). To confirm this, mtDNA copy number was analyzed using qPCR by quantifying mitochondrial ND1 and ND5 relative to the nuclear HK2 gene. A significant reduction in mtDNA copy number was observed in *Dcaf17* knockdown cells (Fig. 2f). Together, these findings suggest that reduced intracellular ROS predominantly arises from a relative reduction in the number of mitochondria within GC-1 spg cells subsequent to the knockdown of *Dcaf17*, underscoring DCAF17’s role in maintaining mitochondrial integrity.

**Figure 2.**
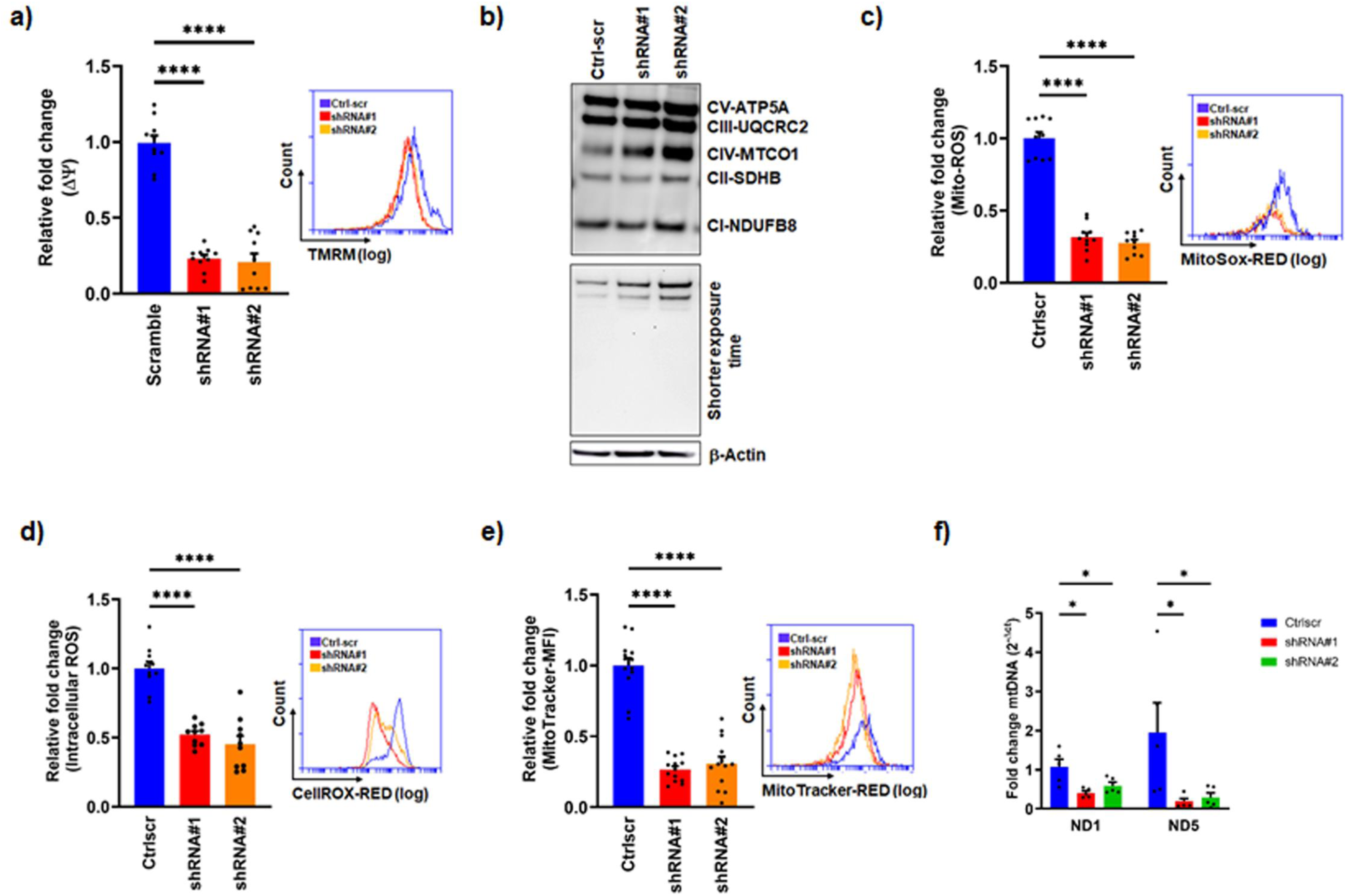
***Dcaf17* depletion disrupts mitochondrial homeostasis and reduces intracellular ROS levels.** (a) Mitochondrial membrane potential (ΔΨm) was measured by TMRM staining and flow cytometry. Fold change relative to scr is shown. Representative histograms from one experiment are included (on the right). (b) Immunoblot analysis of OXPHOS complex proteins in total cell lysates. Lower panel shows the same blot at shorter exposure to visualize low-abundance bands. (c, d) Mitochondrial and total ROS were assessed using MitoSOX Red CMXRos and CellROX Deep Red, respectively. (e) Mitochondrial content was measured using MitoTracker Red staining. (f) Mitochondrial DNA content was assessed by qPCR of ND1 and ND5, normalized to Hk2. Data represent mean ± s.e.m. from at least three biologically independent experiments unless stated otherwise. Statistical analysis was performed using unpaired two-tailed Student’s t-test. p ≤ 0.05 (*), p ≤ 0.01 (**), p ≤ 0.001 (***), and p ≤ 0.0001 (****). Data are shown as AU.

### *Dcaf17* knockdown is associated with ferroptosis induction in GC1 spg cells

Given the observed mitochondrial alterations and previous reports linking Woodhouse-Sakati syndrome (WSS) to increased iron accumulation in the basal ganglia [40], we hypothesize that *Dcaf17*-knockdown may influence iron metabolism in the GC1-spg cells. To investigate whether reduced DCAF17 expression disrupts iron homeostasis, we analyzed the mRNA expression levels of a selected panel of iron metabolism-related genes in *Dcaf17* knockdown cells compared to scramble-cells. The selection of iron metabolism-related genes for expression analysis was based on the panel established by Li-Huang et al. [41], with modifications to better suit our experimental context.

Specifically, four additional genes were incorporated: the glutamate transporter xCT, due to its role in glutathione-mediated thiol-redox regulation and extra-mitochondrial iron–sulfur cluster (ISC) maturation; transferrin (TF), the primary iron-binding transport protein; and mitochondrially encoded cytochrome b (CytB), a critical component of the mitochondrial oxidative phosphorylation (OXPHOS) system. These additions were made to provide a more comprehensive assessment of iron homeostasis and mitochondrial function in *Dcaf17* knockdown cells. RNA extracts derived exclusively from *Dcaf17*-shRNA1-GC-1 spg cells were utilized for the initial screening. Four genes showed significant differential expression in *Dcaf17*-silenced cells (Fig. 3a). Western blot confirmed downregulation of xCT and TF, along with upregulation of transferrin receptor (TFRC). Since TRFC mediates iron uptake, and ferritin (FTH1/FTL1) stores iron, we included both proteins in the analysis. FTH1 was moderately upregulated, while FTL1 expression remained unchanged (Fig. 3b).

**Figure 3.**
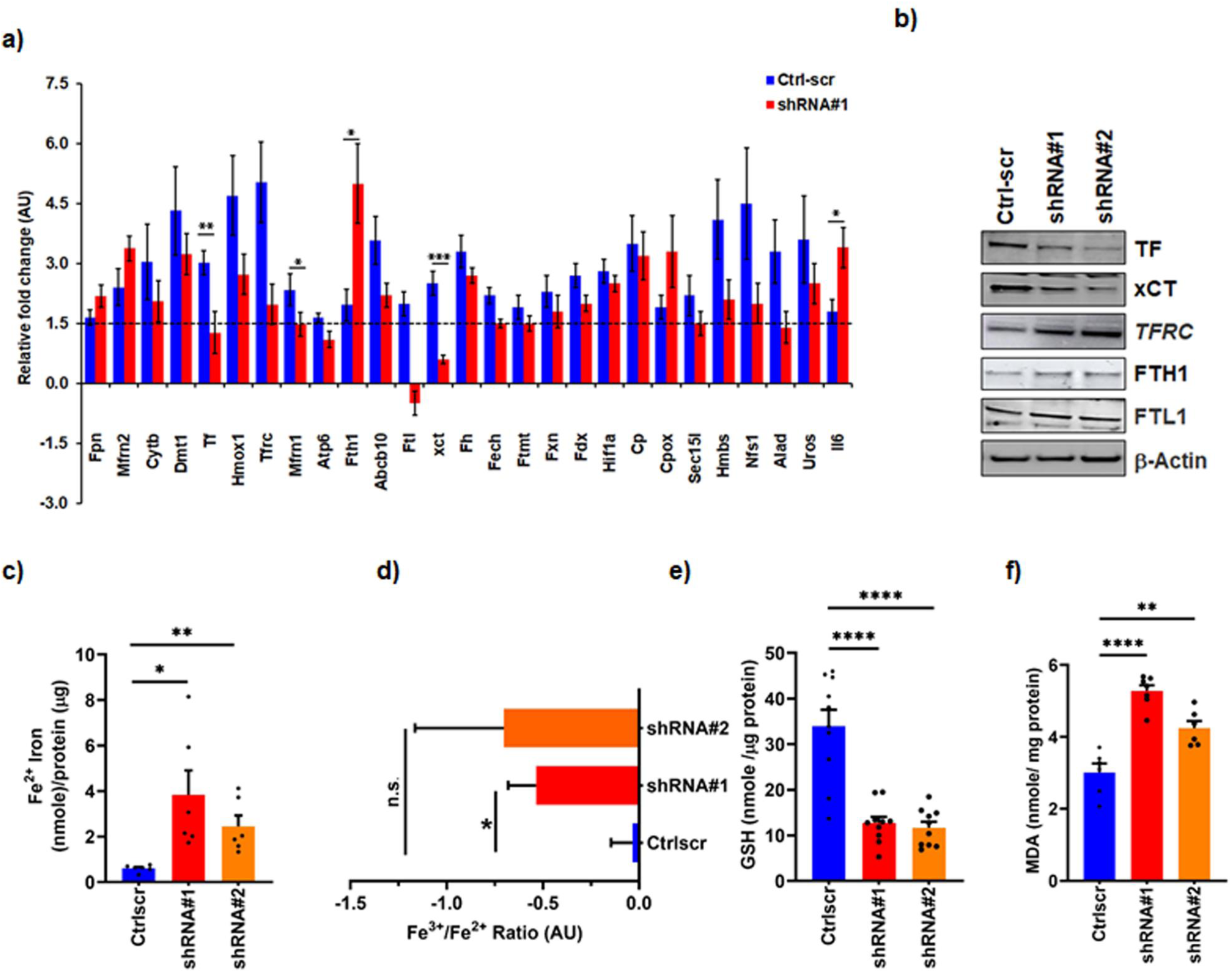
***Dcaf17* knockdown sensitizes GC-1 spg cells to ferroptosis.** (a) Relative mRNA expression of iron metabolism-related genes following *Dcaf17* knockdown (shRNA#1), measured by RT–qPCR. All mRNA levels were normalized to 18S rRNA. (b) Immunoblot analysis of indicated proteins in total cell lysates; β-Actin was used as a loading control. (c, d) Intracellular Fe²⁺ and Fe³⁺/Fe²⁺ ratio was calculated from total and ferric iron measurements. (e) Glutathione (GSH) levels were quantified to assess cellular redox status. (f) Lipid peroxidation was evaluated by measuring malondialdehyde (MDA) levels. Data are presented as mean ± s.e.m. from at least three biologically independent experiments unless otherwise indicated. Statistical significance was assessed using unpaired two-tailed Student’s t-test. p ≤ 0.05 (*), p ≤ 0.01 (**), p ≤ 0.001 (***), and p ≤ 0.0001 (****). Data are expressed as nmol/mg protein or AU.

To determine whether these molecular changes affected intracellular iron levels, we measured ferrous (Fe²⁺) and ferric (Fe³⁺) iron concentrations. Remarkably, *Dcaf17* knockdown significantly increased Fe²⁺ levels and reduced the Fe³⁺/Fe²⁺ ratio (Fig. 3c– d), indicating a shift toward redox-active iron. This iron imbalance is known to promote oxidative damage through lipid peroxidation. Supporting this, knockdown cells showed decreased xCT expression—important for glutathione (GSH) synthesis and ROS defense. We therefore assessed GSH levels and lipid peroxidation. *Dcaf17*-silenced cells exhibited significantly reduced GSH and elevated malondialdehyde (MDA) levels, a marker of lipid peroxidation (Fig. 3 e-f). These results imply that ferroptosis contributes to the cell death observed in *Dcaf17*-deficient spermatogonial cells.

### *Dcaf17* loss impairs lysosomal acidification and function in GC-1 spg cells

Iron is involved in essential biological processes such as DNA biosynthesis and repair, oxygen transport, and energy generation. These cellular functions depend on iron’s capacity to oscillate between its ferrous (Fe^2+^) and ferric (Fe^3+^) forms, and failing to sustain homeostatic levels, which can lead to excessive free Fe^2+^, may cause damage to proteins, nucleic acids, and membrane lipids, ultimately resulting in ferroptosis. Lysosomes are crucial for cellular iron trafficking and metabolism, and the accumulation of iron within lysosomal compartments can compromise lysosomal functionality and cellular homeostasis. Thus, we sought to investigate whether the attenuation of *Dcaf17* expression adversely affects lysosomal acidification and functionality. Lysosomal acidity was assessed using LysoTracker and acridine orange (AO). Both probes showed significantly reduced fluorescence in *Dcaf17*-silenced cells, indicating a perturbation in lysosomal acidity or reduction in lysosome quantity (Fig. 4a-b). To further assess lysosomal function, we measured the protein levels of SQSTM1/p62 using ELISA and immunoblotting. Both assays revealed increased p62 accumulation in *Dcaf17* knockdown cells, consistent with compromised lysosomal functionality (Fig. 4c-d). These results suggest that DCAF17 is essential for maintaining normal lysosomal function and that its loss promotes lysosomal dysfunction.

**Figure 4.**
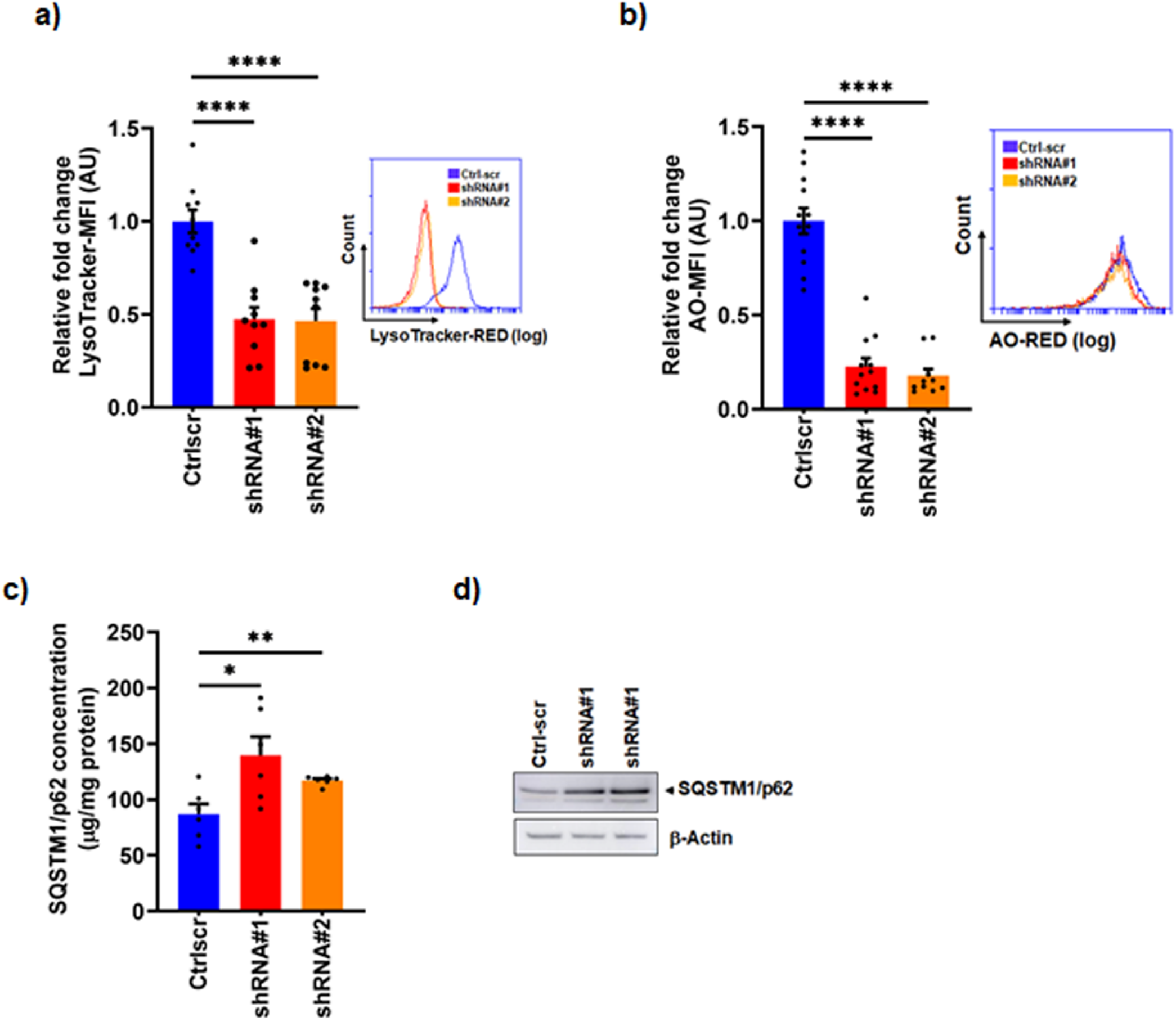
***Dcaf17* knockdown impairs lysosomal function in GC-1 spg cells.** (a) Lysosomal pH was assessed by LysoTracker Red staining and quantified by flow cytometry. (b) Lysosomal membrane integrity was evaluated using acridine orange (AO) staining. Representative histograms from one experiment are included (on the right) (c) SQSTM1/p62 protein levels were quantified by ELISA to evaluate lysosomal degradation activity. (d) Western blot confirmation of p62 accumulation in total cell lysates. Quantification is shown as fold change relative to scr. Data are presented as mean ± s.e.m. from four biologically independent experiments unless otherwise noted. Statistical analysis was performed using unpaired two-tailed Student’s t-test. p ≤ 0.05 (*), p ≤ 0.01 (**), p ≤ 0.001 (***), and p ≤ 0.0001 (****). Data are shown as AU.

### *Dcaf17* knockdown activates necroptotic and inflammatory signaling pathways in GC-1 spg cells

Lysosomal dysfunction plays a pivotal role in inflammation. Moreover, activation of inflammation signaling pathways is closely linked to ferroptosis. Therefore, we assessed the expression of key inflammation-related markers with a focus on those that have been reported to be associated with the chronic inflammatory state within the male reproductive system of murine models [42]. Western blot analysis showed that *Dcaf17* knockdown led to increased levels of JNK1/2, NLRP3, cleaved caspase-1, and cleaved-IL-1b key markers associated with inflammation and necroptosis. Notably, levels of MLKL—a necroptosis executioner—were significantly elevated in *Dcaf17*-silenced cells (Fig. 5b). LDH release, a marker of membrane damage and necroptosis, was also significantly increased (Fig. 5c). The combined findings indicate that the above described increased cell death observed in cells following *Dcaf17* deficiency activates necroptotic and inflammasome-related signaling pathways in GC-1 spg cells.

**Figure 5.**
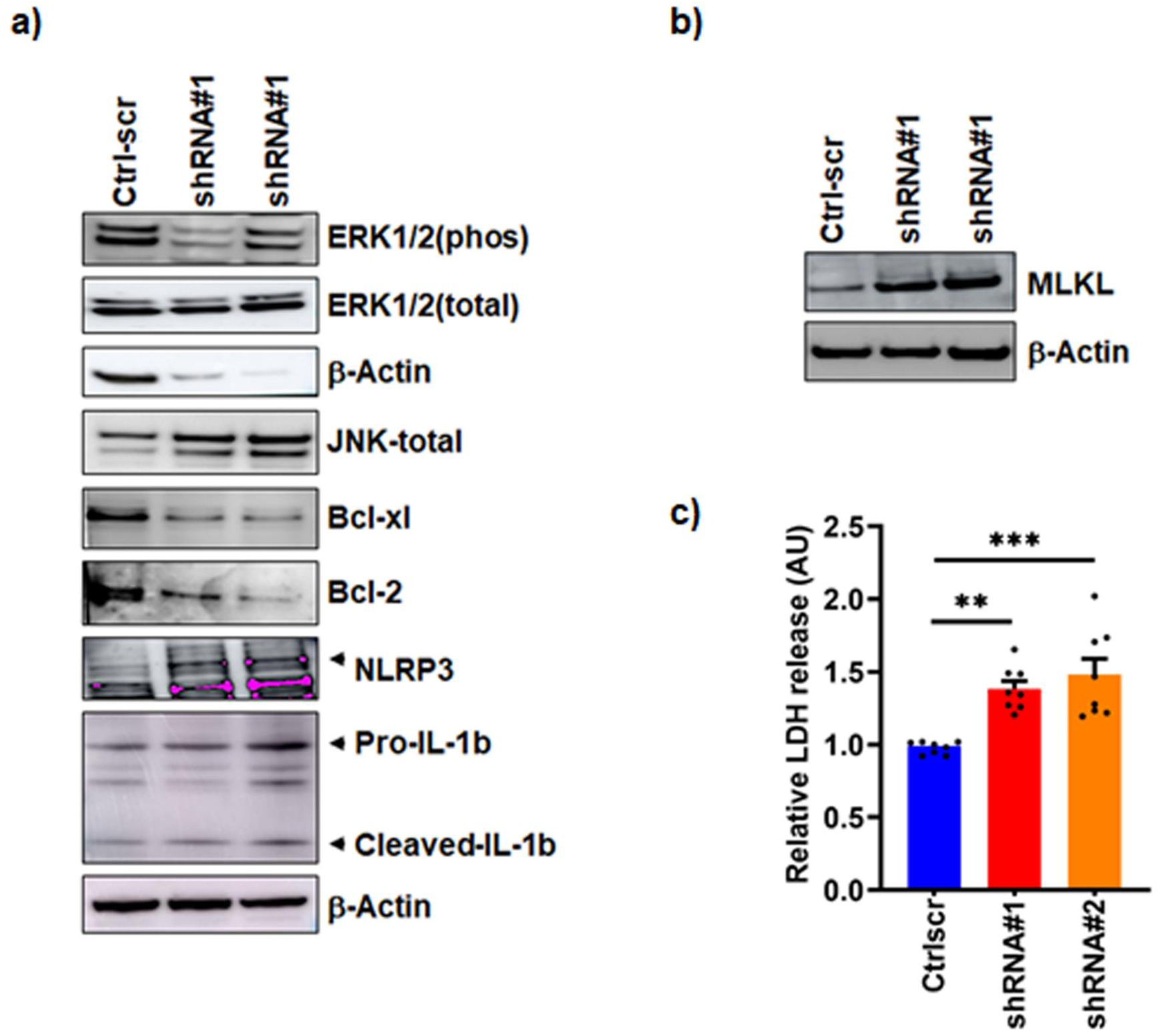
***Dcaf17* knockdown activates inflammatory signaling and promotes necroptosis in GC-1 spg cells.** (a) Immunoblot analysis of total and phosphorylated JNK and ERK1/2, NLRP3, pro-and cleaved IL-1β, and downstream anti-apoptotic markers Bcl-xL and Bcl-2 following *Dcaf17* knockdown. β-Actin served as a loading control. (b, c) Necroptosis was evaluated by MLKL immunoblotting and by measuring lactate dehydrogenase (LDH) release into the medium. LDH is expressed as percentage relative to scr. Immunoblots represent one biological replicate. LDH quantification data are shown as mean ± s.e.m. from at least three independent experiments. Statistical significance was assessed using unpaired two-tailed Student’s t-test. p ≤ 0.05 (*), p ≤ 0.01 (**), p ≤ 0.001 (***), and p ≤ 0.0001 (****). Data are shown as AU.

### *Dcaf17*-related molecular signatures are recapitulated in WSS patient-derived LCLs and *Dcaf17*-KO mouse testis

To assess whether the molecular signatures uncovered in vitro extend to clinically and physiologically relevant contexts, we analyzed lymphoblastoid cell lines (LCLs) derived from three patients with WSS harboring a homozygous *Dcaf17* mutation, alongside testicular tissue and isolated sperm from *Dcaf17*-knockout (KO) mouse. In line with the in vitro results, patient-derived LCLs demonstrated reduced viability (Fig. 6a), decreased mitochondrial membrane potential (Fig. 6b), and diminished mitochondrial mass (Fig. 6c– d). These cells also showed increased Fe²⁺ levels (Fig. 6f), reduced Fe³⁺/Fe²⁺ ratio (Fig. 6e), elevated MDA (Fig. 6g), GSH depletion (Fig. 6h), and lysosomal dysfunction (Fig. 6i– j). In parallel, KO sperm showed a marked decrease in mitochondrial membrane potential (Fig. 7a) accompanied by reduced mitochondrial ROS (Fig. 7b), while KO testes exhibited GSH depletion (Fig.7c), elevated Fe^2+^ iron levels (Fig. 7d) and increased lipid peroxidation (Fig. 7e). Collectively, these findings These consistent findings across in vitro, patient, and mouse models strongly support a conserved role for DCAF17 in maintaining organelle integrity and preventing ferroptosis in male germ cells.

**Figure 6.**
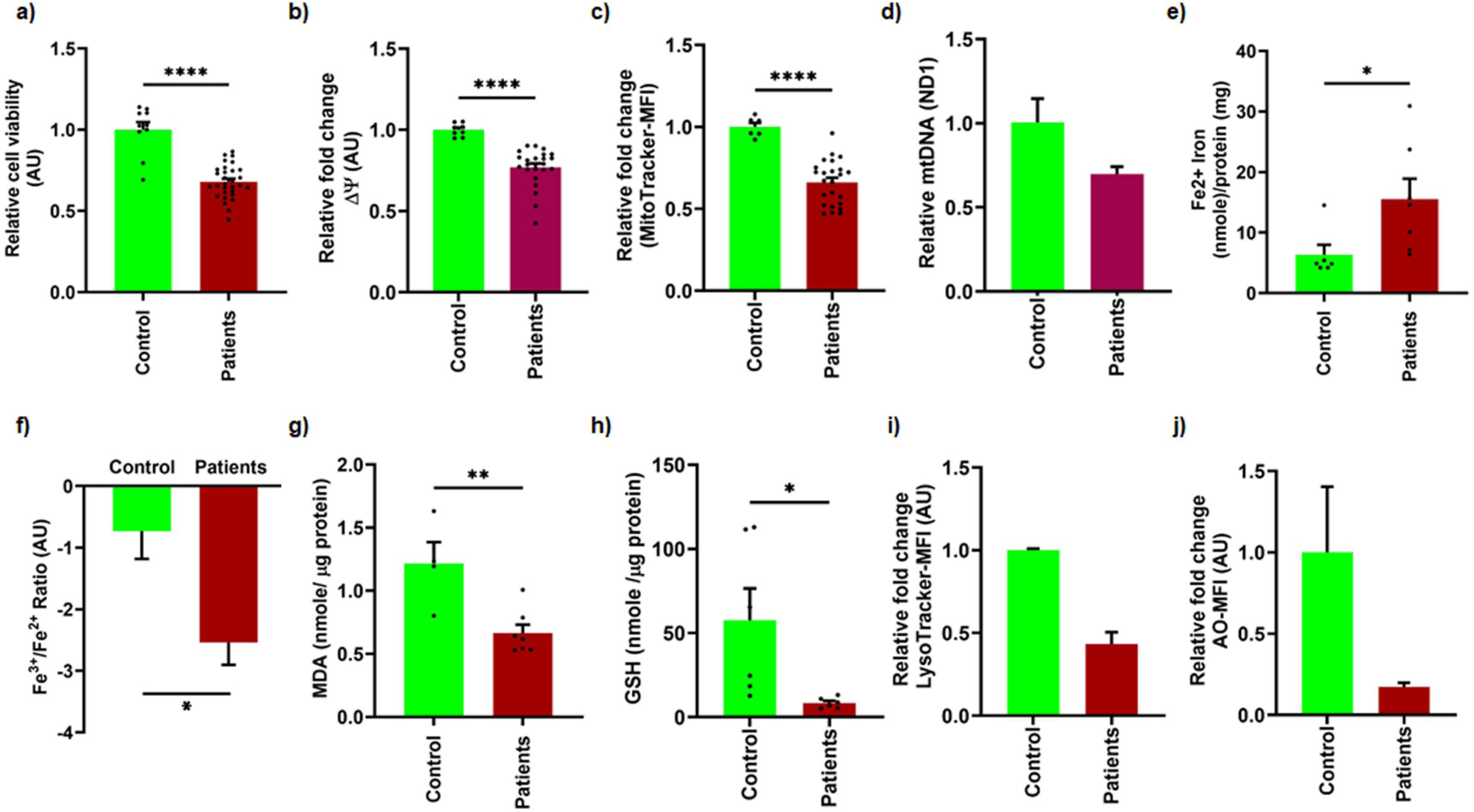
*Dcaf17* depletion recapitulates mitochondrial, lysosomal, and redox dysregulation in patient-derived cells. Cellular phenotypes were evaluated in lymphoblastoid cell lines (LCLs) derived from three WSS patients carrying a pathogenic DCAF17 variant and one control. The following analyses were performed: (a) cell viability, (b) mitochondrial membrane potential, (c–d) mitochondrial mass, (e) intracellular Fe²⁺, (f) Fe³⁺/Fe²⁺ ratio, (g) malondialdehyde (MDA), (h) glutathione (GSH) levels, and (i–j) lysosomal acidification. Data from patient-derived lines were pooled and analyzed as a single biological group. Results are presented as mean ± s.e.m. from at least three biologically independent experiments (with the exception of (i) (n = 2) and (j) (n = 1)). Statistical significance was determined using an unpaired two-tailed Student’s t-test, with p ≤ 0.05 (*), p ≤ 0.01 (**), p ≤ 0.001 (***) and p ≤ 0.0001 (****). Data are shown as arbitrary units (AU).

**Figure 7.**
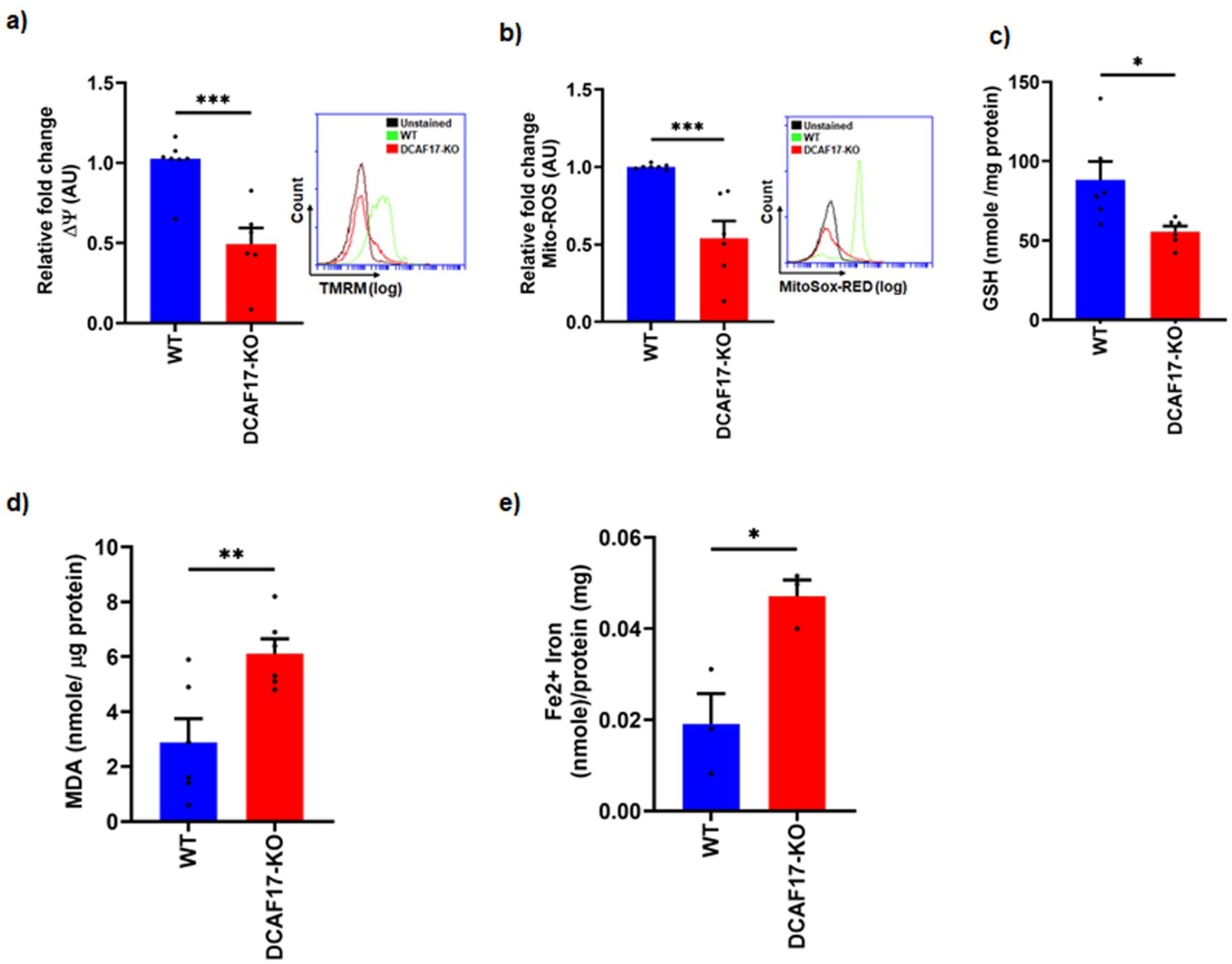
*Dcaf17* depletion induces mitochondrial and redox dysregulation in a knockout mouse model. Cellular and biochemical phenotypes were evaluated in spermatozoa and testicular tissue from *Dcaf17* knockout (KO) mice. (a) Mitochondrial membrane potential and (b) mitochondrial mass were measured in isolated spermatozoa. (c) Glutathione (GSH) and (d) malondialdehyde (MDA), and (e) ferrous iron (Fe²⁺) levels were quantified in testicular tissue. Data represent mean ± s.e.m. from at least four animals per group. Statistical significance was determined using an unpaired two-tailed Student’s t-test with p ≤ 0.05 (*), p ≤ 0.01 (**) and p ≤ 0.001 (***). Data are presented as arbitrary units (AU).

## Discussion

This study uncovers a previously uncharacterized role for DCAF17 in preserving cellular homeostasis and integrity within male germ cells and associated cellular models. Through the use of the GC-1 spg mouse spermatogonia cell line, and subsequent validation in lymphoblastoid cell lines (LCLs) derived from patients with Woodhouse-Sakati Syndrome (WSS) and testicular tissue from *Dcaf17*-KO mice, we identify a cascade of mitochondrial, lysosomal, and ferroptotic dysregulations triggered by *Dcaf17* deficiency. These findings position DCAF17 as a critical factor in preventing iron-dependent cell death and supporting its involvement in male infertility pathogenesis. Silencing *Dcaf17* in GC-1 spg cells resulted in marked reductions in cell viability and increased cell death, highlighting the protein’s essential role in germ cell survival. These findings were consistent with observations in *Dcaf17*-KO mice, which exhibited elevated germ cell apoptosis [34]. *Dcaf17* knockdown resulted in G1 cell cycle arrest in GC-1 spg cells, which model spermatogonia-the male germ cells at the onset of spermatogenesis. Arrest at this stage can disrupt spermatogonial differentiation and ultimately impair fertility [43]. Previous studies have shown that inflammatory mediators such as TNFα and nitric oxide can block cell cycle progression and trigger apoptosis in GC-1 cells [44]. These observations imply that the inflammatory signaling observed in our study may contribute to the G1 arrest.

Mitochondrial dysfunction was evidenced through reduced mitochondrial membrane potential and mass, despite increased OXPHOS protein levels— likely a compensatory response to counteract impaired mitochondrial function [45, 46]. This phenotype is consistent with mtDNA depletion, which disrupts the electron transport chain and limits ATP production, which impairs the electron transport chain and limits ATP production [47]. Interestingly, ROS levels were paradoxically reduced in knockdown cells, consistent with the described effect of mtDNA loss on superoxide generation [48, 49]. As mitochondria are indispensable for spermatogenesis, supporting energy demands, regulating cell fate, and maintaining sperm quality, their dysfunction can lead to impaired sperm development and male infertility [50]. Importantly, mitochondria are central to iron handling through biosynthesis of heme and Fe–S clusters. *Dcaf17* knockdown led to Fe²⁺ accumulation and a decreased Fe³⁺/Fe²⁺ ratio—conditions that promote lipid peroxidation via Fenton chemistry [51]. Iron overload and redox imbalance are hallmarks of ferroptosis and are known to reduce sperm quality and compromise testicular function [52–54].

Iron trafficking and regulation involve intricate interactions between various organelles including mitochondria and lysosomes. The coordinated interplay between these two key organelles is crucial for maintaining cellular iron homeostasis, ensuring that iron is readily available for essential processes while preventing its toxic accumulation. Disruptions in this delicate balance can lead to a cascade of events culminating in iron metabolism dysregulation and cellular dysfunction [55]. Disrupted lysosomal function was indicated by reduced acridine orange and LysoTracker fluorescence and increased SQSTM1/p62 levels. As lysosomes mediate autophagy and intracellular iron recycling, their dysfunction inhibits the clearance of damaged mitochondria and may release labile iron into the cytosol, aggravating oxidative damage. [56, 57]. Mitochondrial defects and lysosomal dysfunction are tightly linked, with impairment in one organelle often leading to dysfunction in the other. While this interdependence has been implicated in cancer progression, neurodegenerative diseases (e.g., Parkinson’s disease), and age-related conditions [4] are now shown to extend to male germ cells. Further mechanistic insight revealed reduced expression of xCT (SLC7A11) and transferrin (TF), alongside upregulation of transferrin receptor (TFRC) and ferritin heavy chain (FTH)-a transcriptional pattern indicative of impaired iron export and glutathione biosynthesis. xCT is important for cystine uptake and glutathione (GSH) synthesis; its loss depletes GSH leading to decreased cellular antioxidant capacity, promoting lipid ROS accumulation and ferroptosis [58]. Increased lipid peroxidation, diminished GSH, elevated Fe²⁺ levels, and reduced Fe³⁺/Fe²⁺ ratios strongly implicate ferroptosis activation as a major cell death mechanism in *Dcaf17*-deficient cells. Although both MitoSOX and CellROX signals were reduced, lipid peroxidation remained elevated, suggesting a shift from mitochondrial to iron-driven, non-mitochondrial ROS generation. This decoupling is consistent with the metal-catalyzed formation of lipid alkoxyl and peroxyl radicals, which are not detected by the superoxide-specific probes used in this study (MitoSOX and CellROX) [59].

Ferroptosis is increasingly recognized as a central mechanism of germ cell loss, with studies highlighting the roles of Fe²⁺ accumulation, mitochondrial dysfunction, antioxidant depletion (e.g., GPX4, SLC7A11, Nrf2), and lipid peroxidation enzymes (e.g., ACSL4, ALOX15) in testicular damage [24, 53, 54, 60, 61]. Our findings extend this framework by identifying DCAF17 as a key regulator of this pathway.

Beyond ferroptosis, *Dcaf17* deficiency also induced necroptosis and NLRP3 inflammasome activation, as evidenced by increased MLKL, NLRP3, cleaved IL-1β, and phosphorylated JNK1/2. Concurrent LDH release and downregulation of Bcl-2 and Bcl-xL further support membrane rupture and inflammation. Since Bcl-2 family proteins also suppress necroptosis, their loss amplifies susceptibility to regulated necrosis [62, 63]. Iron overload, lipid peroxides, and damaged mitochondria have been implicated in NLRP3 inflammasome activation, which amplifies inflammatory signaling and induces pyroptotic cell death [64, 65]. This conjunction of cell death pathways promotes a pro-inflammatory environment detrimental to spermatogenesis [42, 53, 66–68]. Importantly, these molecular signatures were recapitulated in LCLs derived from WSS patients and testicular tissue from *Dcaf17* knockout mice, emphasizing the relevance of these findings to human reproductive biology. In KO mice, sperm exhibited reduced mitochondrial membrane potential and ROS levels, consistent with the in vitro results. These defects likely underlie subfertility or infertility associated with *Dcaf17* deficiency. The cross-model validation strongly suggests that DCAF17 serves as a master regulator of redox balance, organelle integrity, and cell fate in germ cells.

Ferroptosis and NLRP3 activation have been linked to idiopathic and environmentally induced infertility. Clinical studies reported reduced GPX4 and xCT expression, along with elevated ROS, MDA, and iron levels in testicular conditions such as varicocele, diabetes, hepatitis, and non-obstructive azoospermia [24, 53, 54, 60, 61]. Environmental toxins are also known to promote ferroptosis in Sertoli, Leydig, and germ cells by disturbing iron homeostasis and antioxidant defense mechanisms [53]. Additionally, NLRP3 is upregulated in sterile testicular inflammation, implicating it in immune-mediated impaired spermatogenesis [67, 68].

This study is the first to demonstrate that *Dcaf17* deficiency initiates a cascade of mitochondrial and lysosomal dysfunction, iron dysregulation, ferroptosis, necroptosis, and inflammasome activation, emphasizing its crucial role in maintaining male reproductive health. Moreover, these findings offer new insights into the pathophysiological mechanisms underlying WSS syndrome. Further investigation is warranted to delineate the tissue-and context-specific pathways through which DCAF17 orchestrates cellular homeostasis.

## Acknowledgments

We would like to thank Mr. Mohammed Rajab and Mrs. Maha Alanazi for their valuable assistance with mouse handling, genotyping, and tissue collection. Their support was instrumental in the successful completion of the in vivo studies.

